# Lipid rafts are the new Stress Granules regulators

**DOI:** 10.1101/2025.03.04.641411

**Authors:** Anaïs Aulas, Coralie Di Scala

## Abstract

Stress granules are cytoplasmic inclusions^1^ with cyto-protective functions^2-6^ assembling in response to stress. They are now accepted to be part of the pathological mechanism in several diseases, from cancer to neurodegenerative disorders^7-10^. However, the field is still struggling to find common regulators of their assembly and function^7,11^.

In this study, we describe an unraveled mechanism involving lipid raft, via gangliosides and cholesterol, in the regulation of SG formation. This is the first report about regulation of SG by the cell membrane composition. This discovery could have a significant impact on the understanding of several disease mechanism.

**MATERIAL AND METHODES:** *Cell culture & cell treatment:* MDA-MB-231 (ATCC) and SH-SY5Y (ATCC) cells were maintained at 37 °C with 5% CO_2_ in Gibco Dulbecco’s Modified Eagle Medium: Nutrient Mixture F12 (DMEM-F12, GIBCO, Waltham, MA, USA) supplemented with 10% Fetal Bovine Serum (FBS, Eurobio, Les Ulis, France), 20 mM HEPES (GIBCO, Waltham, MA, USA), 1X Penicillin streptomycin (GIBCO, Waltham, MA, USA). Cells are treated with methyl-β-cyclodextrine (MβCD) (MDA-MD-231 5mM, SH-SY5Y 1mM) 48h before experimentation, or with d,l-threo-l-Phenyl-2-hexadecanoylamino-3-morpholino-1-propanol (PPMP) (MDA-MD-231 5μM, SH-SY5Y 10μM) for 24h.

*Immunofluorescence:* Cells were seeded on coverslips, treated 48h with PPMP or 24h with MβCD before the experiment. After stress treatment, cells are washed quickly with PBS before to be fixed for 15min with 4% Paraformaldehyde (Thermo Scientific, Waltham, MA, USA) in PBS. Cells were then permeabilized and blocked with IF buffer PBS-0.3% TX100 (Euromedex, Souffelweyersheim, France), 1% Glycine (Sigma, Saint-Louis, MO, USA), 5% Normal Horse Serum (Sigma, Saint-Louis, MO, USA), 5% Bovine Serum Albumine (Sigma, Saint-Louis, MO, USA) for 30 min at room temperature. Primary antibodies (**Table S1**) were diluted in IF buffer and incubated 1 h at room temperature. Coverslips were washed three times for 5 min with 1X PBS between primary and secondary antibody incubations. Subsequently, secondary antibodies (**Table S1**) were added along with DAPI for 1 h at room temperature in IF buffer. Cells were washed extensively 3 times with 1X PBS and mounted with ProLong Antifade reagent (Invitrogen, Carlsbad, CA, USA). Pictures were taken with confocal microscope LEICA LSM880

*Western Blot:* Following drug(s) treatment(s), cells were washed with phosphate-buffered saline (PBS) and lysed in RIPA buffer (150mM NaCl, 50mM Tris pH7.4, 1%TritonX100, 0.1% SDS, 1% Sodiun desoxycholate) with Halt phosphatase and protease inhibitors (Thermo Scientific). Laemmli’s sample buffer supplemented was added to samples to 1X final concentration. Samples were boiled, 5min 95°C before being loaded on a NuPAGE™ 4–12% Bis-Tris gel (Invitrogen) and transferred to nitrocellulose membrane (GE Healthcare). Membranes were blocked with Tris-buffered saline with 0.1% Tween-20 (TBS-T) with 5% BSA for at least 30 min at room temperature. Antibodies were diluted in 2.5% BSA in TBS-T. Primary antibodies were incubated overnight at 4°C and secondary antibodies for 1 h at room temperature; mouse anti G3BP1 antibody (Santa Cruz sc-365338), rabbit anti Caprin-1 antibody (ProteinTech Group 15112-1-AP), mouse anti puromycin antibody (Millipore MABE342), mouse anti GAPDH (abcam ab8245). Antibody detection was performed using SuperSignal West Pico Chemiluminescent Substrate (Thermo Scientific). Revelation of the blot was made using G:BOX machine (Syngene) via the GeneSys software. Blot analysis and quantification were done using ImageJ software.

*Statistical Analysis:* Statistical analyses were done on 3 independent experiments. Student T-TEST were performed to compare control to PPMP samples or control to MβCD samples.

## INTRODUCTION

Stress granules (SG) are cytoplasmic inclusions that assemble after cells exposition to stress^1,12^. Those inclusions composed of proteins and mRNA^1,10^ help the cells to overcome stress exposition^2-6^. Indeed, stress exposition induces a global translation repression and directs untranslated mRNA into SG^13,14^. This will 1) protect from stress induced degradation^2^ and 2) give the translation priority to stress responsive proteins such as chaperones^13^. This action saves energy for the cells^15^ and allows the efficient translation restart as soon as stress is removed^2^. On top, SG assembly protects cells from cell death by inhibiting the action of pro-death protein via their recruitment into the structure^4-6,8^. These properties prompt researches to investigate the link between SG and human diseases and SG are now related to several human disorders from neurodegenerative diseases^10^ to cancer^8,9^. The control of SG assembly is mainly studied by up or down regulation of proteins or by finding new drugs for their induction or inhibition^7,11^.

The SG regulation in diseases is still under intensive investigation and is directing the research to unexplored diseases-induced dysregulations. Among them, the contribution of lipid rafts in those processes have been overlooked. This is a gap since lipids field have unraveled dysregulation in virtually all human pathologies. More specifically, profound lipid alterations have been observed in cancer and neurodegenerative diseases and are now known as key features of these diseases^16-19^. Additionally, lipid rafts regulate cell signaling in neuronal cells^20^ and cancer cells^21^. This evidence that membranes lipids composition can interfere with cell signaling. In this study we investigate the potential effect of well-known lipids rafts perturbing drugs of the cell ability to assemble SG.

## RESULTS

We chose to investigate the potential regulation of SG by membrane lipids using two cell lines: MDA-MB-231 (breast cancer cell line) and SH-SY5Y (neuroblastoma cell line). We subjected both cell lines to an increased concentration of sodium arsenite (SA), a cell stress inductor, to setup a baseline of sensitivity. To follow SG we follow G3BP1 and Caprin-1, two specific SG markers that colocalize in cytoplasmic foci ^22^ (**Fig. 1A&B**). Each cell line has its own sensitivity, since SH-SY5Y cells start assembling SG at a lower SA concentration (25 μM) than MDA-MB-231 cells (50 μM) (**Fig. 1A&B**). Increasing SA concentration enhances the proportion of SG positive cells in both cell line until reaching the maximum at 100 μM for MDA-MB-231 (97,5±0,6%) and 50μM for SH-SY5Y (93,4±4,2%).

**Figure 1:**
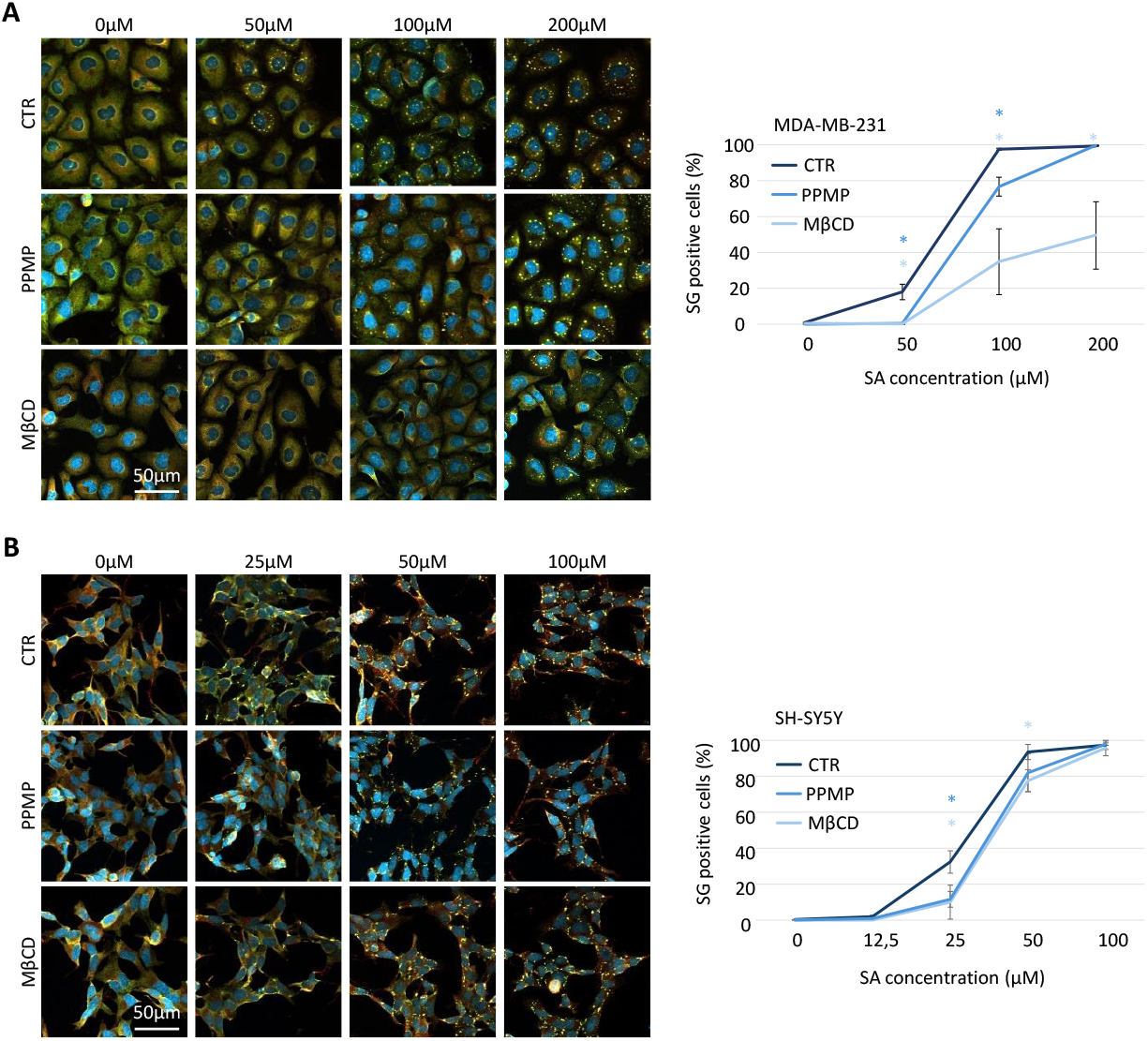
Cells with lack of gangliosides and cholesterol at the membrane required higher dose of SA to induce the assembly of SGs. **A-** MDA-MB-231 cells are treated with PPMP (5μM, 48h) or MβCD (5 mM, 24h) before SG experimentation. The day of the experimentation cells were treated 1h with SA at the indicated concentration and collected. **B-** SH-SY5Y cells are treated with PPMP (10 μM, 48h) or MβCD (1 mM, 24h) before SG experimentation. The day of the experimentation cells were treated 1h with SA at the indicated concentration and collected. **A&B-** After collection cells are then fixed and stained with G3BP1 (green), Caprin-1 (red) for SG labelling and DAPI (bleu) for nuclei visualization before to by imaged using confocal microscopy. Left: representative pictures. Right: quantifications. N=3, *p<0,05

Then to investigate the role of lipid rafts in SG regulation we took advantage of methyl-β-cyclodextrine (MβCD) that removes cholesterol from the plasma membrane and d,l-threo-l-Phenyl-2-hexadecanoylamino-3-morpholino-1-propanol (PPMP) that interferes in the ganglioside synthesis pathway. Under both treatments, cells had reduced ability to assemble SG regardless cell lines (**Fig. 1A&B**). PPMP and MβCD decrease similarly the cell ability to induce SG in SH-SY5Y (PPMP –20,8±2,1% at 25 μM; MβCD –22,3±5,5% at 25 μM and –15,9±5,2% at 50 μM). No difference for either treatment under 100μM SA was observed. On MDA-MB-231 PPMP decreases the number of SG positive cells under 50 and 100 μM SA to 13,7±2,3% and 22,6±7,5% respectively. Whereas MβCD decreases drastically the cell ability to assemble SG (−20,0±5,3, 34,9±18,3 and 49,5±18,8% at respectively 50, 100 and 200μM SA).

The cells with altered lipid rafts composition on cholesterol and gangliosides were less likely to assemble SG. We also investigated if altered lipid rafts composition was able to delay the assembly of SG at the highest SA concentration. We quantified SG assembly after 15-, 30– and 60-min exposition to 200 µM SA for MDA-MB-231 cell. On basal condition, where lipid rafts are not altered, SG assembly started after 15 min of SA exposition (1.9±0.4%) and rapidly increased over time (30 min, 79.6±6.5% 60 min 96.8±.08%) (**Fig. 2A**). SG assembly slowed down when lipid rafts are perturbed. When cholesterol is removed under MBCD treatment, no SG are observed after 15 min exposition to 200 µM SA and only 20.1±10.4% and 57.8±28.8% of cells assembled SG after respectively 30 and 60 min to SA 200 µM (**Fig. 2A**). When gangliosides pathway is altered, the SG assembly kinetic also slowed down with 0.6±0.6% of SG positive cells after 15 min exposition to 200 µM SA, 52.8±16.5 after 30 min and 83.6±3.6% after 60 min exposition to 200 µM SA (**Fig. 2A**). For SH-SY5Y cells the assembly does not start before 30min (data not shown) and follow the same trend as the MDA-MB-231 cells. PPMP delays the assembly of SG of 45,7±11,5 and 29,7±12,1 and MβCD of 33,3±7,4% and 29,9±3,4% after an exposition of 30 and 45min to 100 μM SA respectively **(Fig. 2B)**.

**Figure 2:**
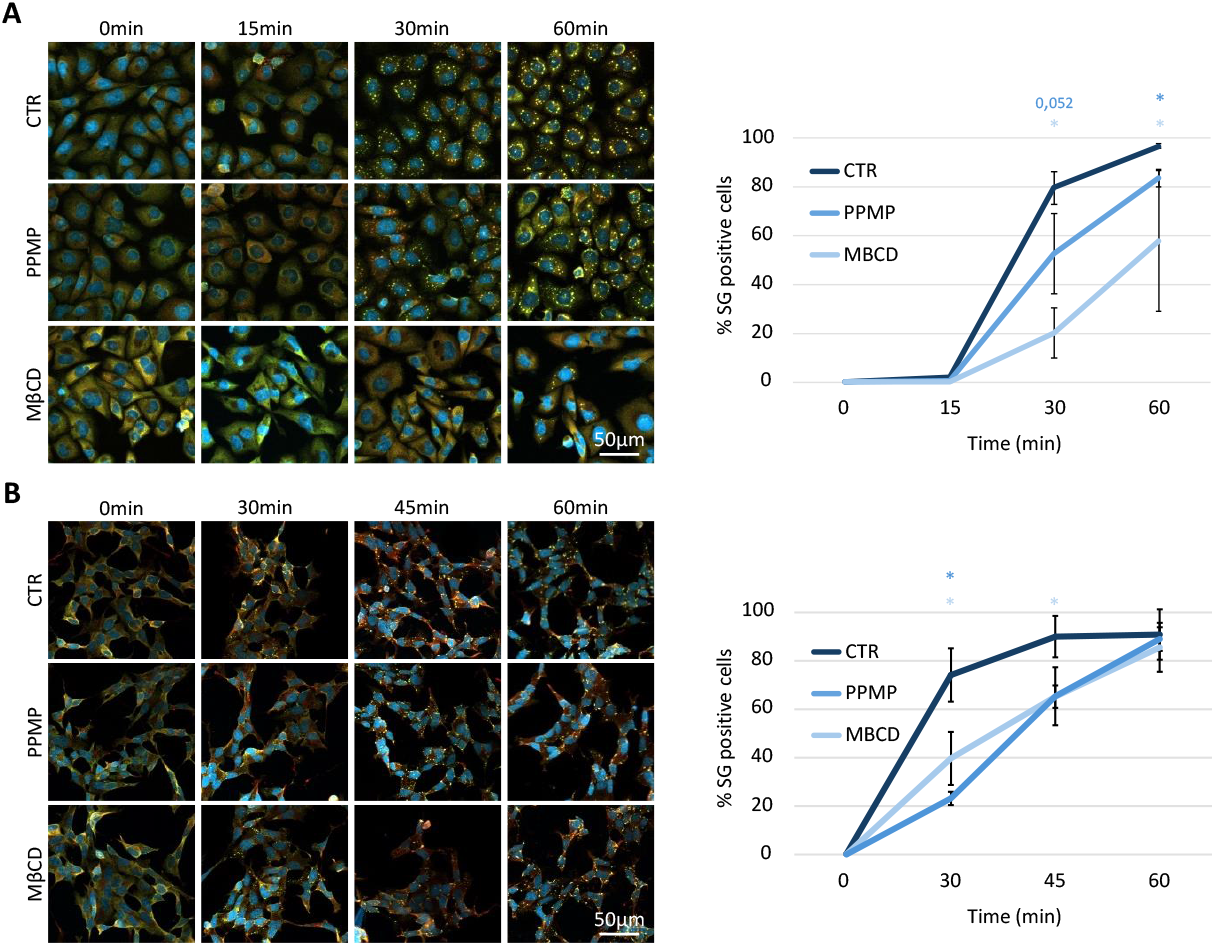
Lack of gangliosides and cholesterol at the membrane delay the formation of SGs. **A-** MDA-MB-231 cells are treated with PPMP (5 μM, 48h) or MβCD (5 mM, 24h) before SG experimentation. The day of the experimentation cells were treated with 200μM SA and collected at the indicated time. **B-** SH-SY5Y cells are treated with PPMP (10 μM, 48h) or MβCD (1 mM, 24h) before SG experimentation. The day of the experimentation cells were treated with 100 μM SA and collected at the indicated time. A&B-After collection cells are then fixed and stained with G3BP1 (green), Caprin-1 (red) and DAPI (bleu) before to by imaged using confocal microscopy. Left: representative pictures. Right: quantifications. N=3, *p<0,05

To ensure that foci observed under PPMP and MβCD are bona fide SG and not unspecific aggregagtion^12^, we subjected cells to puromycin that enhances SG formation by releasing mRNA into the cytoplasm^23,24^ or Cycloheximide (CHX) that inhibits SG formation by trapping mRNA in the ribosome^24,25^ (see **Fig. 3A** for more detailed explanation on the mechanisms of these drugs). Without addition of SA neither the puromycin nor the CHX induce the assembly of SG in any of our samples (**Fig. 3B**). Control, PPMP or MβCD treated cells assemble SG under 100 μM SA, and this assembly is inhibited by the CHX (**Fig. 3C**). Meanwhile, SG formation is enhanced by the puromycin treatment in all cells treated with 50 μM SA (**Fig. 3D**). The cell response to puromycin and CHX treatment on top of the double labelling with G3BP1 and Caprin-1 refers to the formation of bona find SG in cells depleted in ganglioside or cholesterol at the membrane.

**Figure 3:**
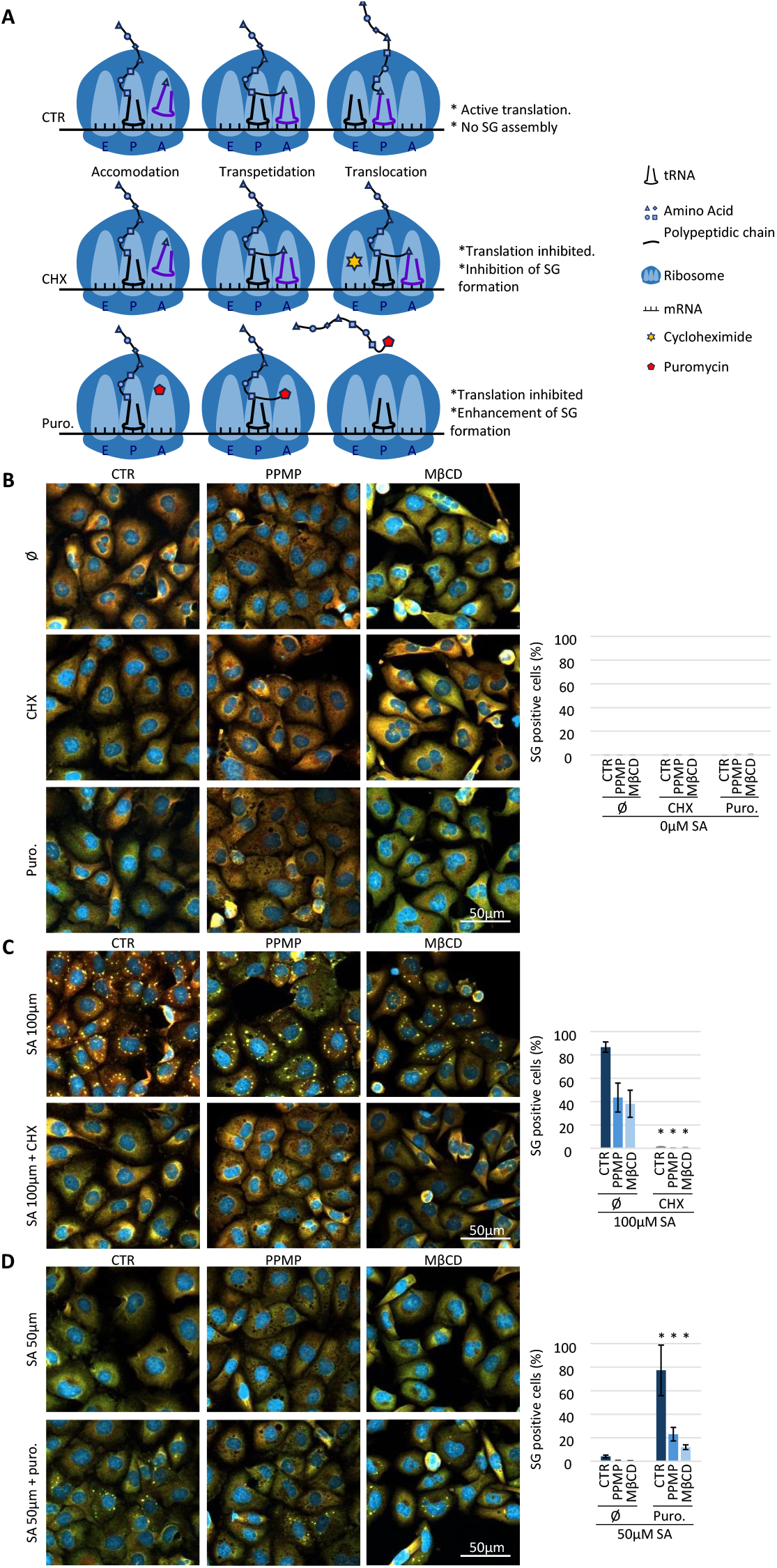
Foci induced after lipid raft disruption are SGs. **A-** Mechanism of action on translation of CHX and puromycin. CTR) Translation in basal condition: 1. tRNA binds in the A binding site (acceptor). 2. Peptide bond then forms between the new amino acid and the forming peptide chain in the P (Polypeptide) site.3. The tRNAs translocate to the E (Exit) site, where the first tRNA can exit the ribosome. CHX) Inhibition of translation by cycloheximide: When cycloheximide is present, it binds to the E site of the ribosome. As a result, the translocation step cannot take place and translation stops with the RNAs being translated blocked in the ribosome. Puro.) Inhibition of translation by puromycin: When cells are treated with puromycin, the latter integrates with nascent polypeptide chains, stopping translation and releasing mRNAs and polypeptide chains into the cytoplasm. **B-D-** MDA-MB-231 are treated with PPMP (5 μM, 48h) or MβCD (5 mM, 24h) before SG experimentation. SG experimentation is the indicated combination of CHX (50 μM) or Puromycine (20 μM) and/or Sodium Arsenite (50 μM or 100 μM as indicated) for 1h.Ø: no CHX, no puromycine. CHX: Cycloheximide. Puro.: Puromycin. SA: Sodium Arsenite. Cells are then fixed and stained with G3BP1 (green), Caprin-1 (red) and DAPI (bleu) before to by imaged using confocal microscopy. Left: representative pictures. Right: quantifications. N=3, *p<0,05 **B-** Cells are treated with CHX, puromycin and compared to untreated samples (Ø) without any SA addition. **C-** All samples are stressed using 100μM of SA to induce a robust SG response (Ø). CHX is added to inhibit SG formation. **D-** Cells are treated with sub-optimal stress (50 μM) to induce minimum or no SG (Ø). Puromycin is added to enhance SG formation.

SG formation is a well-regulated process. Currently there are two major pathways that could regulate the formation of SG; by avoiding the translation repression^26^ or by affecting the level of expression of the level of expression of the SG core protein G3BP1/2^3,11^ (**Fig. 4A**). We choose to investigate both options. First we investigate the translation repression happening following stress exposure by using the puromycylation technic^27^. Cells are pulse 5min with low concentration of puromycin that will incorporate into nascent polypeptidic chain (**Fig. 3B&C**). Puromycin detection using a specific antibody will be then representative of active translation. For the control sample the translation repression occurs already at 50 μM SA, whereas under PPMP and MβCD treatment a higher dose of SA (100 μM) is required to induce translation repression compared to the pre-stressed sample. On top, treatment with PPMP and MβCD reduces the G3BP1 global level of expression (**Fig. 3B&D**.

**Figure 4:**
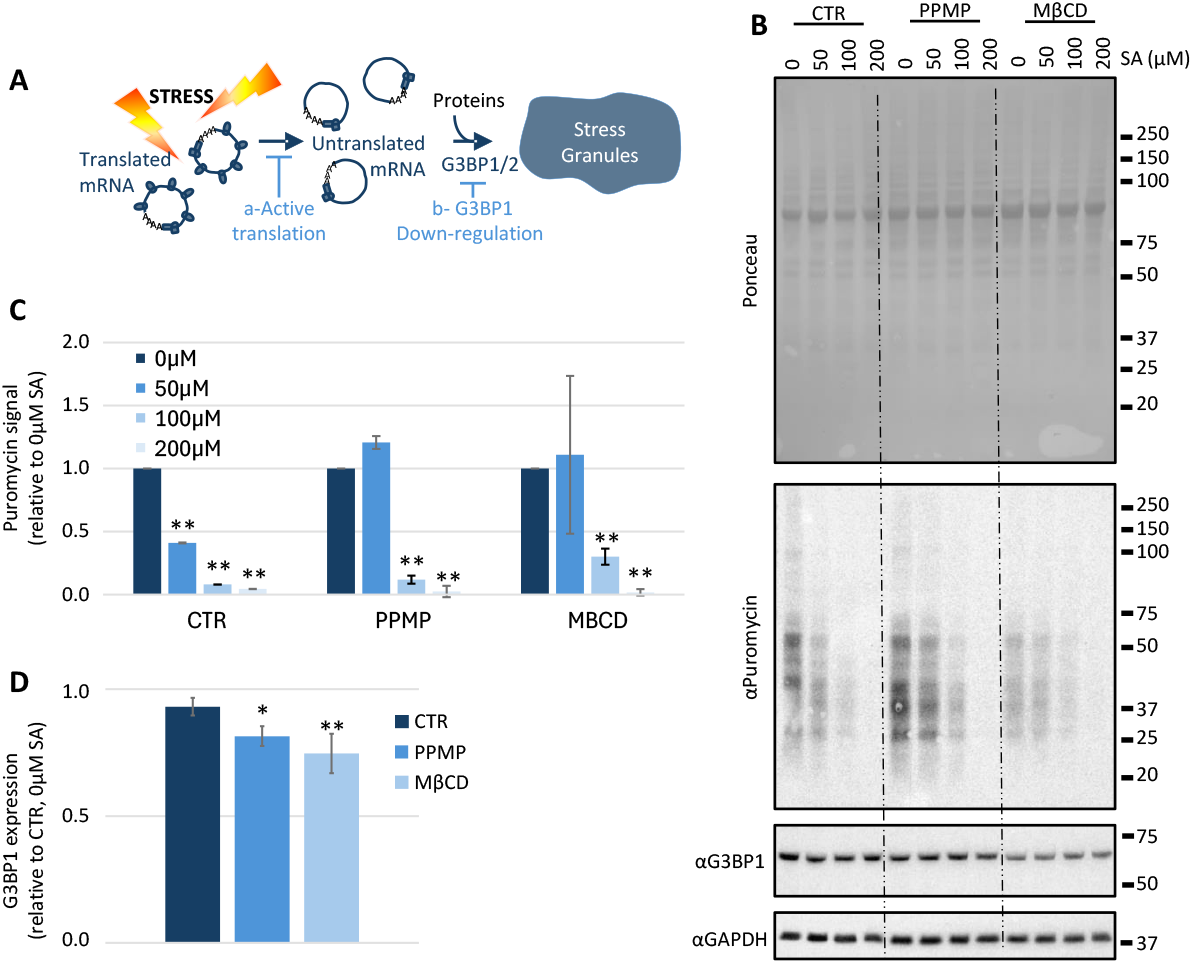
Modification of lipids raft decrease G3BP1 expression level and translation inhibition in response to stress. **A-** Under stress exposure, polysome disassemble to allow mRNA to be recruited to SG with proteins. ***a-*** Keeping the translation active will prevent the assembly of SG. ***b-*** The downregulation of one of the two scaffolding protein G3BP1 or G3BP2 will also (partially) prevent the formation of SG. **B-D-** MDA-MB-231 MDA-MB-231 are treated with PPMP (5μM, 48h) or MβCD (5mM, 24h) before to be exposed to the indicated SA concentration for 1h. 5min before cell lysis, cells were pulsed with puromycin 5μg/ml. N=3, *p<0,05, **p<0,001 **B-** Representative blots. **C-D-** Quantifications. Signal intensity was measured using ImageJ software. Each band intensity was expression relatively to the GAPDH intensity of the same sample and then expression was plotted relatively to the untreated sample (CTR, 0μM SA). **C-** Puromycin level, representative of general protein expression. **D-** G3BP1 expression. All the CTR, PPMP and MBCD treated samples were respectively pulled together for analysis.

## DISCUSSION

Currently the SG field is still intensively investigating the link between SG and dysregulated biological process^28^. Especially how some proteins mutations could affect the assembly and or function of SG in human pathogenesis. Recent studies show that lipids dysregulation is of growing interest in the development of human diseases^16-19^ prompting us to investigate the role of these kind of regulation in the assembly of SG.

Here we found that cholesterol and gangliosides are key molecules for the fast and efficient SG formation. Removing cholesterol and decreasing gangliosides levels in plasma membrane of cells reduces the expression level of G3BP1, one of the key scaffolding proteins for SG assembly^11^. On top they also reduce the ability of the cell to induce translation inhibition, the starting point for the SG assembly cascade^13,29^. The combination of the two leads to the delay in the formation of SG and reduces sensitivity of cells to stress. As SG are known as pro-survival entities to help cell to overcome stress exposition/insult^4-6,8,30^, we speculate that cells via the dysregulation of their lipid composition within the plasma membrane may become more vulnerable to stress exposition. This is particularly relevant in the context of neurodegenerative disorders where SG assembly is reduced, and lipid composition of neuronal cells is profoundly modified ^31,32^(ref). On the other hands the over-representation of cholesterol^33,34^ together with change of gangliosides nature and level in cancer cells^17^ could increase the ability of cells to answer to stress exposition and provide pro-survival properties to those cells.

Even if a link between SG formation and lipids droplet formation had already been proven^35^, the implication of lipid membrane has never been investigated before. This study put in light that lipids dysregulation could have an underestimated impact on pro-survival mechanism and therefore being directly involved in human pathogenesis through SG regulation. The link between SG and lipids will need further investigation to elucidate the exact mechanism in action and how to counteract them. This study opens promising perspectives.

## AUTHOR CONTRIBUTIONS

Original idea: AA & CDS. Conception and realization of the experiments and figures: AA. Writing: AA & CDS. Funding: CDS

## ACKOWLEDGMENT

This work has been supported by the Academy of Finland (grants 333096 and 335956) to CDS. We also thank the HiLIFE fellow program and the Neuroscience Center (University of Helsinki) who supported CDS as a young group leader.

